# Synergid calcium ion oscillations define a new feature of pollen tube reception critical for blocking interspecific hybridization

**DOI:** 10.1101/2020.07.23.218263

**Authors:** Nathaniel Ponvert, Mark A. Johnson

## Abstract

Reproductive isolation leads to the evolution of distinct new species, however, the molecular mechanisms that promote and maintain reproductive barriers are elusive. In flowering plants sperm cells are immotile and are delivered to female gametes by the pollen grain which germinates a polarized extension, the pollen tube, into floral tissue. After growing via polar extension to the female gametophyte, the pollen tube signals with receptive female cells flanking the egg in two distinct steps. If signaling is successful, the pollen tube releases sperm cells for fusion with female gametes. To better understand cell-cell recognition during reproduction, we investigated calcium ion dynamics associated with interspecific cell signaling between mating partners. We observed that interspecific pollen tubes successfully complete initial cell-cell signaling, but fail during later phases of pollen tube reception. Our work refines our understanding of pollen tube reception as a critical block to interspecific hybridization. Our results also shed light on the functional significance of the two phases of pollen tube reception, and implicate the second step as being the stage during which the pollen tube presents its genetic identity.

## Introduction

Pollen tube reception is one of the final potential pre-zygotic barriers to interspecific hybridization. At high rates during interspecific crosses, sperm are not released and reproduction is unsuccessful^1,2^. Loss of function mutations in female gametophyte-expressed signaling components FERONIA/LORELEI and NORTIA, that participate in both the first^1,3^ and second^4^ stages of pollen tube reception^5^, respectively, result in pollen tube reception failure, concurrent with a phenotype similar to what is observed during interspecific pollination where the pollen tube fails to release sperm. Previous studies using genetically-encoded calcium ion sensors found that contact between the pollen tube tip and the female gametophyte initiates calcium ion oscillations in the synergid cells of the female gametophyte^5,23-25^ in a FERONIA/LORELEI-dependent manner^5^. These oscillations persist for approximately 45 minutes, during the first phase of pollen tube reception. The second phase of pollen tube reception follows the cessation of the initial calcium ion oscillations after which a brief rise in cytoplasmic calcium ion concentration occurs, shortly before pollen tube burst and sperm release^5^.

Male factors also play roles in pollen tube reception, as loss of pollen tube-expressed MYB transcription factors results in defective pollen tube reception as well^6-8^. Given the similarity in sperm release defects between interspecific crosses and losses of function in female and male genes, we took a confocal imaging-based approach to uncover how pollen tube reception plays a role during presentation of species identity. We used a female gametophyte-expressed genetically-encoded calcium ion sensor to monitor the progression of signaling steps during pollen tube reception between interspecific pollen tubes and *feronia* and *lorelei* mutant lines. We also used *myb97,101,120* triple-mutant pollen for crosses with the *feronia* and *lorelei* mutants. Strikingly, we found that interspecific pollen tubes elicit small molecule signaling responses in synergid cells of other species in a FERONIA/LORELEI-dependent manner, as was observed during same-species interactions. Our results suggest that genetic identity is communicated downstream of FERONIA/LORELEI activity. We also observed that *myb* triple-mutant pollen tube suitors elicited small molecule responses in female cells despite an ultimate failure to release sperm. These results are more similar to what we observed during interspecific pollination than to the lack of oscillations observed during crosses onto *lorelei* or *feronia* null mutants, suggesting that MYB-dependent genes act downstream of FERONIA/LORELEI and may define a portion of the pollen tube’s genetic identity.

## Results

The species concept includes the idea that members of different species cannot successfully mate with one another. Interspecific hybridization can fail after a zygote is produced, however, in many systems pre-zygotic barriers prevent zygote formation^1,2,9,10^. Active mechanisms that block pollen tube growth of exotic species have been described in the *Solanaceae, Eucalyptaceae*, and *Brassicaceae*^11-14^. Additionally, molecular incongruity resulting from divergence of proteins that mediate gamete or gametophyte attraction or reception^1,2,15^ can also result in pre-zygotic barriers to interspecific hybridization. The molecular recognition events that determine cellular compatibility between gametes of the same species are only now beginning to be understood^15-21^. The final step in flowering plant fertilization, sperm and egg fusion, is not likely to be species-specific because it is mediated by the highly conserved HAP2 protein, which has been shown to be functionally interchangeable between distantly related species^22^. Therefore, it is not known which male or female signaling components fail during incongruous interspecific crosses with otherwise compatible mating partners, or even when exactly during mating partner interactions the communication of genetic identity fails in these cases when there is no active block to interspecific hybridization. To address these questions, we used well characterized plant models, including *Arabidopsis thaliana*, in the *Brassicaceae* family.

### Same-species pollen tubes induce characteristic synergid cell calcium ion oscillations

To begin our analysis of interspecific interactions, we first analyzed interactions between same-species pollen tubes and synergid cells, as had been done previously^5,23-25^ (Figure 1, Supplementary Figure 1a, Supplementary Video 1). Although two phases of calcium oscillations in synergids were identified previously^5,23-25^, their functional significance remains unknown. *Arabidopsis thaliana lorelei* or *feronia* loss-of-function mutants fail to receive same-species pollen tubes, resulting in a characteristic pollen tube overgrowth phenotype similar to that observed during interspecific pollination^1-3,15^. In *lorelei* and *feronia* mutant females, pollen tube reception defects are accompanied by a severe attenuation in calcium ion oscillations in the cytoplasm of the synergid cells, indicating that *lorelei* and *feronia* mutant synergid cells are oblivious to the arrival of the pollen tube tip^5^ (replicated here in Figure 1c,d,e, Supplementary Figure 1b-c, Supplementary Videos 2,3). To directly measure changes in calcium ion oscillations, we calculated the height of every peak identified in each calcium ion trace. When we compared the average height of maxima identified in calcium ion traces from wild-type synergids to the height of maxima identified in calcium ion traces from *lorelei* and *feronia* mutant synergid cells receiving same species wild-type pollen tubes, we found that in either mutant background there is a statistically significant reduction in average calcium ion oscillation height, which is consistent with previously published results^5^ (Figure 1e).

**Figure 1:**
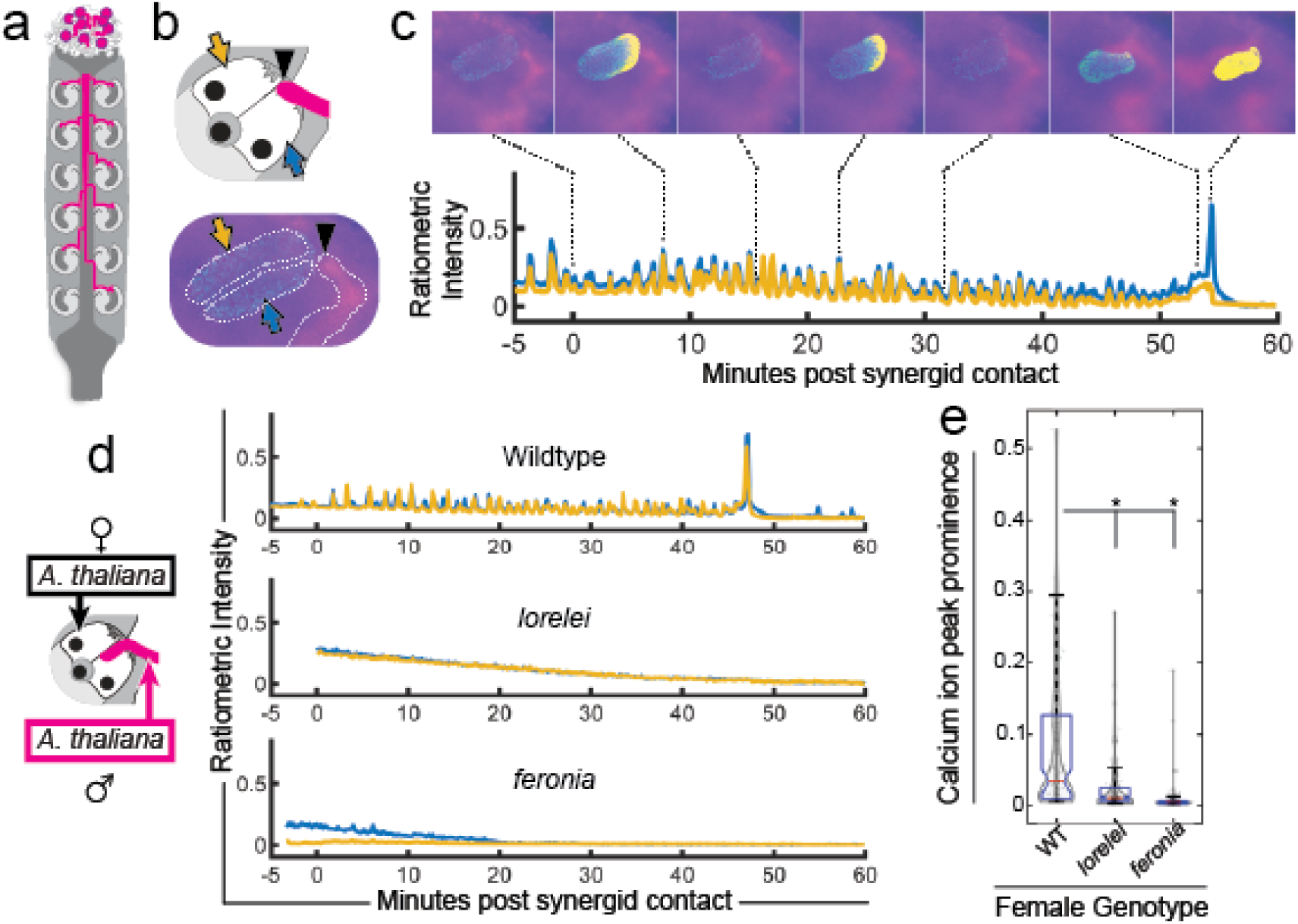
Live imaging of calcium ion oscillations during pollen tube reception. **a.** Pollen tube growth begins after the pollen grain lands on the stigmatic tissue. **b.** The pollen tube makes contact with synergid cells after growing into the embryo sac. Pollen tube tip denoted with black arrowhead, persistent synergid (relative to pollen tube tip) marked with yellow arrow, receptive synergid marked with blue arrow. **c.** The pollen tube tip participates in a multi-step cell signaling event with the receptive synergid cell and induces calcium ion oscillations in the cytoplasm of the synergid cells. **d.** Example traces (traces with mean peak prominence are presented as examples; all other traces are presented in the supplement) from crosses onto wildtype, *lorelei*, and *feronia* mutant females using wildtype *Arabidopsis thaliana* pollen. Calcium ion trace from left synergid cell plotted in yellow and right synergid cell plotted in blue. **e.** Plot of calcium transient prominence observed during crosses onto female genotypes (wildtype n=10 traces n=128 peaks, *lorelei* n=10 traces n=161 peaks, *feronia* n=10 traces n=118 peaks). Statistical significance calculated using analysis of variance (ANOVA, α = 0.05).

### Interspecific pollen tubes induce calcium ion oscillations in synergid cells of different species

We next asked whether calcium ion oscillations occur in synergid cells during interspecific crosses using *Arabidopsis thaliana* females receiving pollen tubes from *Arabidopsis lyrata* or *Olimarabidopsis pumilia* and if the nature of the oscillation is tied to specific recognition. We chose *A. lyrata* and *O. pumilia* as pollen donors, as they are predicted to be diverged from *A. thaliana* ∼4-10 million years ago (MYA) and ∼10-14 MYA, respectively, thereby serving as representatives of closely- and more distantly-related species^26-28^.

When we performed crosses using pollen from either *Arabidopsis lyrata* (n = 10) or *Olimarabidopsis pumilia* (n = 10), we observed the initiation of calcium ion oscillations in wild-type *Arabidopsis thaliana* synergid cells (Figure 2a,c, Supplementary Figure 2a, Supplementary Figure 3a, Supplementary Videos 4,7). However, the calcium ion oscillations induced by either type of interspecific pollen tube (Figure 2) did not appear to follow the stereotypical pattern of progression observed during same-species crosses described above (Figure 1)^5,23-25^. During interspecific interactions, pollen tubes were able to induce calcium ion oscillations in synergid cells, similarly to what is observed during phase one of same-species pollen tube reception, but we very rarely observed a progression into the second phase of reception (a shift from calcium ion oscillation to constitutive increase in calcium ion concentration, termed ‘plateau’) and we did not observe pollen tube burst in any interspecific cross, indicating that induction of calcium ion oscillations alone is not sufficient to induce pollen tube burst. These results suggest that *A. lyrata* and *O. pumilia* pollen tubes are able to successfully signal their arrival to a gametophyte of a different species, but that successful sperm release requires congruity in a distinct molecular recognition event that follows initiation of calcium oscillations.

**Figure 2.**
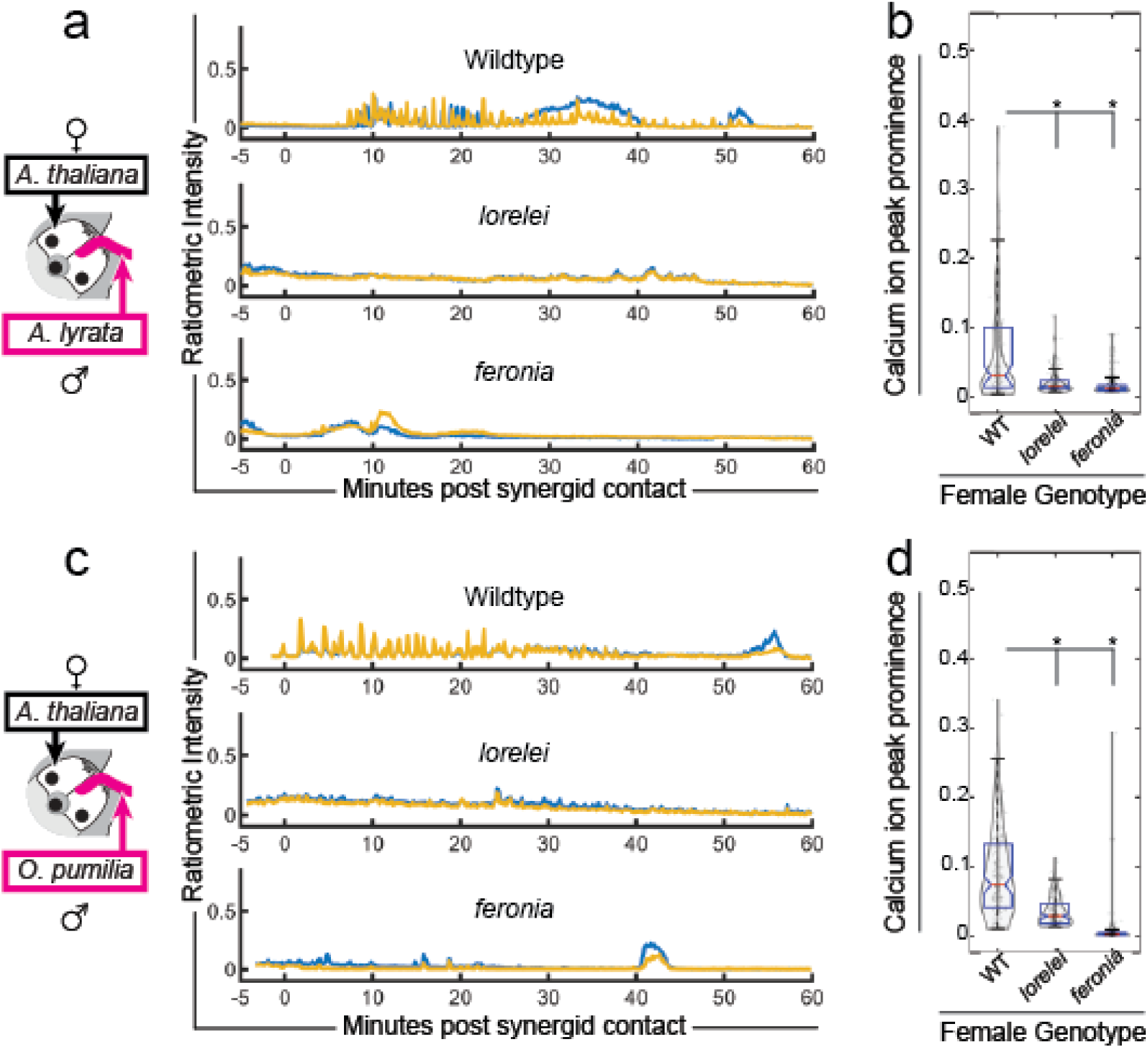
Interspecific pollen tubes elicit calcium ion oscillations in wildtype synergid cells. **a.** Example traces from crosses onto wildtype, *lorelei*, and *feronia* mutant females using wildtype *Arabidopsis lyrata* pollen. **b.** Plot of calcium transient prominence observed during crosses onto female genotypes (wildtype n=10 traces n=82 peaks, *lorelei* n=10 traces n=96 peaks, *feronia* n=10 traces n=121 peaks). Statistical significance calculated using analysis of variance (ANOVA, α = 0.05). **c:** Example traces from crosses onto wildtype, *lorelei*, and *feronia* mutant females using wildtype *Olimarabidopsis pumilia* pollen. Calcium ion trace from left synergid cell plotted in yellow and right synergid cell plotted in blue. **d.** Plot of calcium transient prominence observed during crosses onto female genotypes (wildtype n=10 traces n=90 peaks, *lorelei* n=10 traces n=121 peaks, *feronia* n=10 traces n=81 peaks). Statistical significance calculated using analysis of variance (ANOVA, α = 0.05).

### Calcium ion oscillations induced by interspecific pollen tubes require the FERONIA/LORELEI receptor complex

To investigate whether the calcium ion oscillations observed during interspecific crosses are FERONIA/LORELEI signaling complex-dependent, we crossed *A. lyrata* or *O. pumilia* pollen onto *A. thaliana lorelei* or *feronia* females. If the calcium ion oscillations observed during interspecific pollinations are also dependent on FERONIA/LORELEI as they are during same-species pollination, then an attenuation of calcium ion responses as seen in same-species pollination events would be expected^5^ (Figure 1d,e). Indeed, we observed an attenuation of calcium ion oscillations in *lorelei* and *feronia* mutant females receiving interspecific pollen tubes (Figure 2a-d, Supplementary Figure 2b-c, Supplementary Figure 3b-c, Supplementary Videos 5,6,8,9), similar to those previously reported for *lorelei* and *feronia* mutants receiving wildtype *A. thaliana* pollen tubes^5^ (replicated in Figure 1, Supplementary Figure 1b-c, Supplementary Videos 2,3). These results suggested that FERONIA and LORELEI are required for the interspecific pollen tube-induced calcium ion oscillations but the failure to present congruous identification signals to the receptive synergid cell occurs downstream of FERONIA/LORELEI activation, and a failure of this downstream step results in failure to release sperm.

### *myb97,101,120* transcription factor triple-mutant male pollen tubes phenocopy interspecific signaling defects

We next explored the possibility that the pollen tube is the source of signaling molecules that underlie the molecular recognition of the pollen tube by the synergid cell. To date, the only male-specific mutations resulting in failure of sperm release are triple null mutants in the pollen tube-expressed MYB97, 101, and 120 transcription factors^2,6-8,29^. The *myb97,101,120* triple mutant displays a characteristic pollen tube overgrowth phenotype after making contact with the female gametophyte that is similar to the phenotype observed with *feronia* and *lorelei* mutant females, and to what is observed during interspecific crosses^1-3,6-8,15,29^. However, it is not known when MYB-dependent genes act during reception, and/or if they interact with FERONIA/LORELEI signaling complex. Given the similarity in pollen tube overgrowth phenotypes observed during interspecific pollinations and *myb97,101,120* mutant pollinations, we hypothesized that the MYB-dependent gene regulatory network could constitute some portion of the genetic identity that must be properly perceived by the female gametophyte during pollen tube reception.

To more precisely define the roles of male factors in pollen tube reception, we used *myb97,101,120* triple mutants as pollen donors in same-species crosses. When *myb97,101,120* pollen was crossed to wild-type *A. thaliana* pistils, pollen tubes elicited aberrant calcium ion oscillations within wild-type synergid cells (Figure 3a, Supplementary Figure 3a, Supplementary Video 10). These results supported the possibility that *myb97,101,120* pollen tubes only fail during the second phase of pollen tube reception, similar to interspecific pollinations, and that the MYB-dependent genes could define some portion of the pollen tube’s genetic identity that is presented to the female gametophyte.

**Figure 3.**
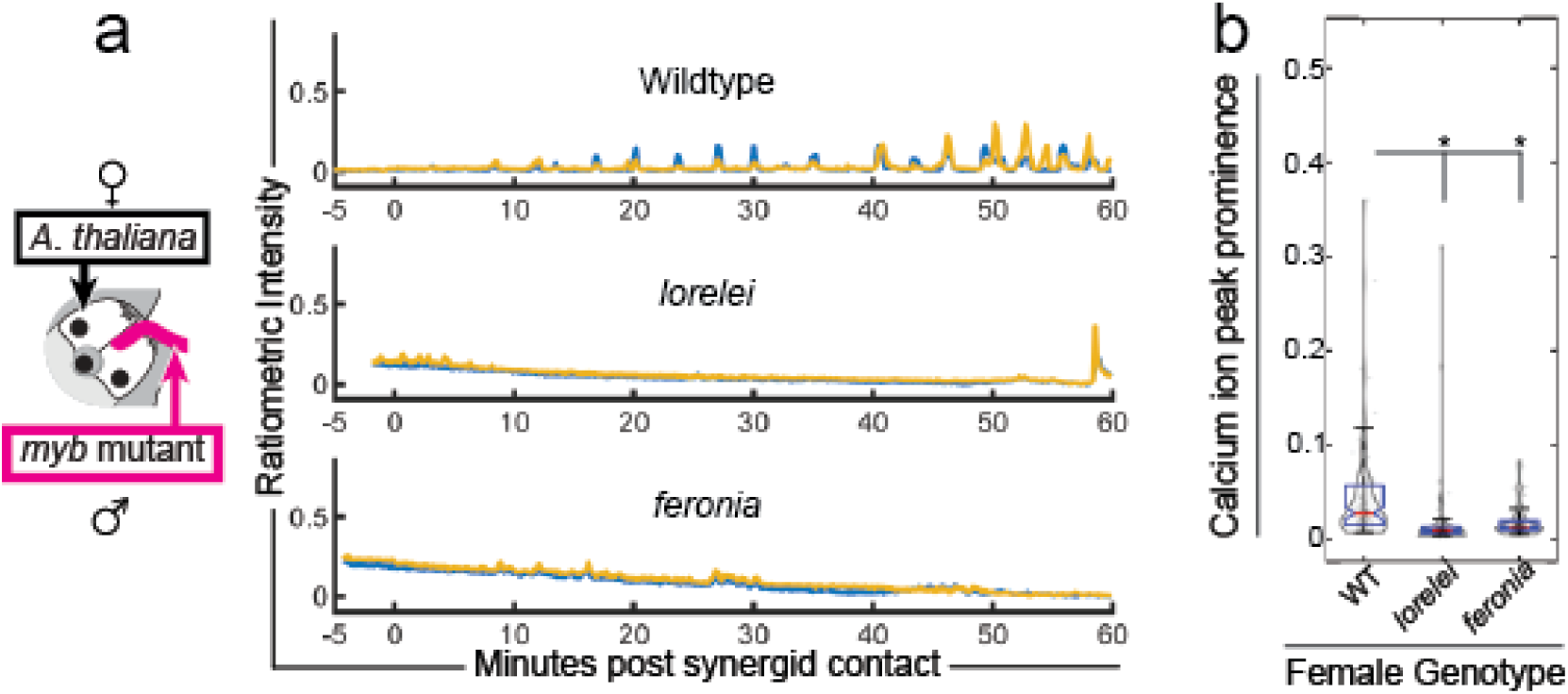
*myb 97, 101, 120* pollen tubes elicit calcium ion oscillations lacking a second phase in wildtype synergid cells. **a.** Example traces from crosses onto wildtype, *lorelei*, and *feronia* mutant females using *myb97,101,120* triple-mutant *Arabidopsis thaliana* pollen. Calcium ion trace from left synergid cell plotted in yellow and right synergid cell plotted in blue. **b.** Plot of calcium transient prominence observed during crosses onto female genotypes (wildtype n=10 traces n=115 peaks, *lorelei* n=10 traces n=137 peaks, *feronia* n=10 traces n=103 peaks). Statistical significance calculated using analysis of variance (ANOVA, α = 0.05).

### Calcium ion oscillations induced by *myb* mutant tubes require FERONIA/LORELEI

Induction of calcium ion oscillations by *myb97, 101, 120* pollen tubes in wild-type synergid cells suggested that the role of MYB factors in pollen tube reception is downstream of FERONIA/LORELEI. To test this possibility, we performed same species crosses between *myb* triple mutant pollen on *lorelei* and *feronia* mutant females. We observed that, as in crosses with wild-type and interspecific pollen tubes, calcium ion oscillations were severely attenuated in *lorelei* and *feronia* mutant females receiving *myb97,101,120* mutant pollen tubes (Figure 3a,b, Supplementary Figure 3b-c, Supplementary Videos 11,12). In the majority of crosses (8/10 in *lorelei* x *myb* and 6/10 in *feronia* x *myb*) calcium transients are largely ablated, with the remaining fraction displaying an occasional calcium ion transient. In such cases, calcium ion transients persist briefly and are, on average, less prominent than in crosses onto wildtype females (Figure 3b). These results supported the conclusion that the MYB gene regulatory network plays a role downstream of FERONIA/LORELEI, during the second phase of pollen tube reception when mate genetic identity is vetted by the female gametophyte.

### Calcium ion oscillation frequency is not the sole determining factor of pollen tube burst success rate

Next, we asked whether the calcium ion oscillation signature recorded from wild-type *Arabidopsis thaliana* synergid cells responding to same-species, interspecific, and *myb* mutant pollen tubes differs. Previous studies have found that cell-cell signaling between roots and fungal or bacterial symbionts initiates calcium ion oscillations in plant root cells, and propose that the frequency of the oscillations depends on the species identity of the symbiont^30-32^. When we measured the frequency of the calcium ion oscillations induced by pollen tubes in our system, we found that *Arabidopsis lyrata* and *Olimarabidopsis pumilia* tubes induce oscillations with similar frequency to those induced by same-species pollen tubes. However, the calcium ion oscillations induced by *myb*-mutant pollen tubes occur with a lower frequency than those induced by wild-type *Arabidopsis thaliana* pollen tubes (Figure 4, Supplementary Figure 5). This suggests that pollen tubes lacing the MYB97,101,120 gene regulatory network are defective in communicating with synergid cells during the second phase of pollen tube reception. However, we concluded that while the frequency of the calcium ion oscillations induced by interspecific and mutant pollen tubes is variable, we did not observe a connection between the frequency of the calcium ion oscillations and interspecific identity.

**Figure 4.**
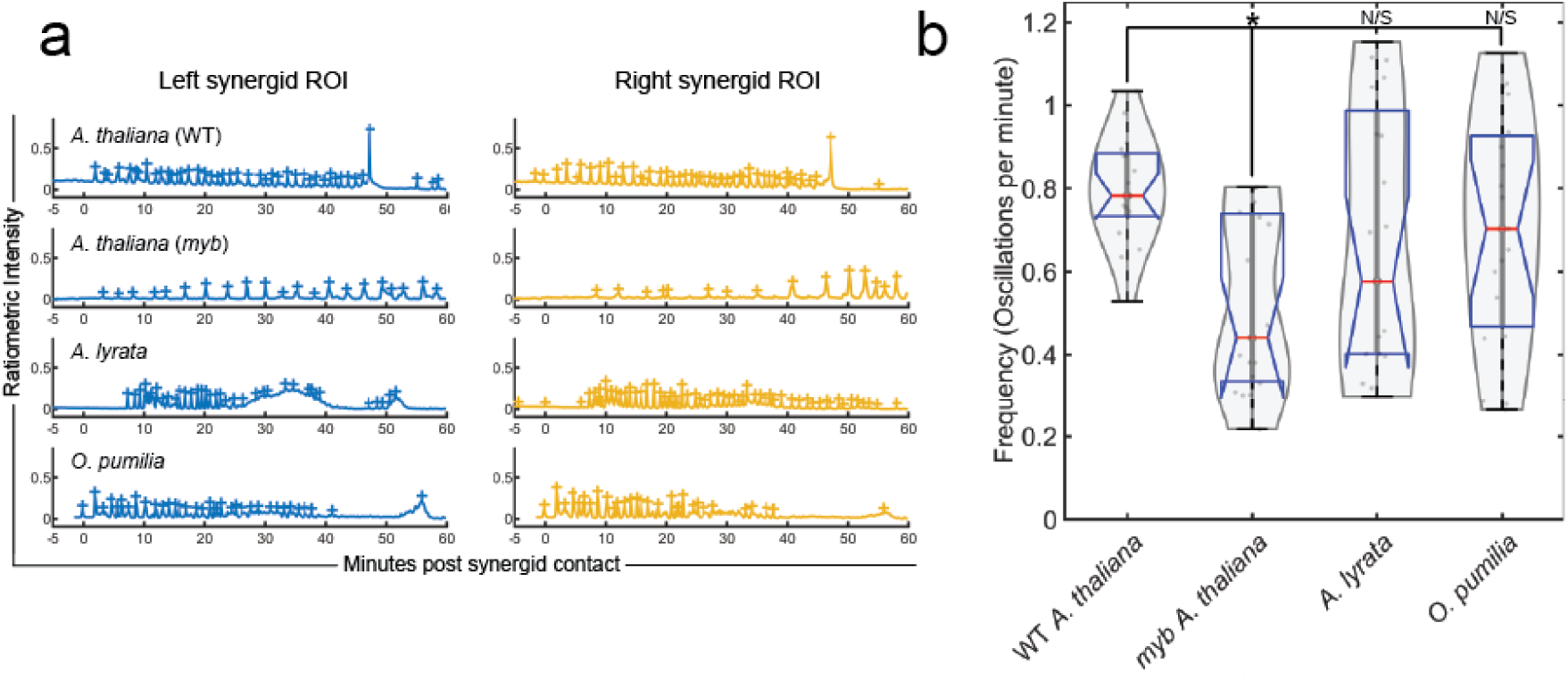
Calcium ion oscillation frequency is lower in synergid cells interacting with *myb* mutant pollen tubes, but not in synergid cells interacting with interspecific pollen tubes. **a.** Frequency of calcium ion oscillations was calculated for wild-type *Arabidopsis thaliana* synergid cells responding to wild-type same-species pollen tubes, *myb* triple-mutant same-species pollen tubes, *Arabidopsis lyrata*, and *Olimarabidopsis pumilia* pollen tubes was calculated by dividing left- and right-synergid ROIs and applying thresholding based on oscillation prominence. Oscillations above the threshold of 5x the minimum prominence observed in the trace are marked with ‘+’. The median replicate is shown. **b.** *myb* triple-mutant pollen tubes induce calcium ion oscillations that occur with lower frequency than wild-type same-species induced oscillations, but interspecific pollen tubes induced oscillations with a frequency not significantly different from same-species pollen tubes. Statistical significance calculated using analysis of variance (ANOVA, α = 0.05).

## Discussion

Here we report on calcium ion oscillations that are induced in the cytoplasm of synergid cells after the arrival of pollen tubes of either an exotic species (*A. lyrata* or *O. pumila*) or same-species (wild type or *myb97,101,120* triple mutant). We found that in both cases the initiation of calcium ion oscillations was dependent on the FERONIA/LORELEI signaling complex. During crosses with the *myb* mutant, calcium ion oscillations occurred with a lower frequency than during wild-type crosses, but interspecific pollen tubes had no significant effect on calcium ion oscillation frequency, suggesting that the frequency of calcium ion oscillations may not be indicative of genetic identity in this system. Our results suggest that pollen tube reception defects (overgrowth and failure to release sperm manifested as a pollen tube coiling phenotype) during interspecific crosses, and during crosses using *myb97,101,120* mutant pollen is not due to a failure in FERONIA/LORELEI function, but rather that after FERONIA and LORELEI recognize the arrival of the pollen tube tip, there is a defect in pollen tube reception that occurs during the second phase. Our findings suggest that the function of the second phase of pollen tube reception is to scrutinize the genetic identity of the pollen tube and present a barrier to interspecific pollen tubes. Additionally, our findings indicate that the MYB97, 101, 120 gene regulatory network may define some portion of the molecular species identity that is presented to the female gametophyte. Finally, our results suggest that the factor or factors that the FERONIA/LORELEI signaling complex is recognizing may be independent from MYB regulation.

It is unclear at this point what factors the FERONIA/LORELEI signaling complex is responding to in the case of pollen tube reception. In sporophytic tissue it is known that FERONIA is capable of binding and responding to Rapid Alkalinization Factor (RALF) peptides^33-35^. It has also been established that pollen tube expressed paralogs of FERONIA interact with pollen tube- and ovule-expressed RALFs that control pollen tube cell wall integrity (pollen tube burst and sperm release) in conjunction with pollen tube expressed-LORELEI paralogs^36-38^. However, no pollen tube-expressed RALF has been identified that interacts with FERONIA/LORELEI directly. It may be the case that physical deformation of the synergid cell surface by the growing pollen tube tip is sufficient to initiate the FERONIA signaling cascade as it is in seedlings^39^, or that pollen-expressed ligands (e.g. RALF peptides) are unaffected by *myb* mutations.

Previous reports^1^ have suggested that FERONIA may be involved in scrutinizing the pollen tube’s genetic identity, but our results suggest that species recognition occurs downstream of the activation of this receptor complex. Using the calcium ion oscillation signature as a readout for points of failure during pollen tube reception has allowed us to order the signaling components and genetic identity factors in this signaling pathway. In plants, signaling between the pollen tube and cells of the pistil and female gametophyte provide an opportunity for investigating molecular interactions determining interspecific mating success. Pollen tube reception (sperm release) breaks down in crosses between closely related species and we have shown that pollen tube-induced and FERONIA/LORELEI-dependent calcium ion oscillations occur in interspecific crosses but are insufficient to induce sperm release. Future work will be aimed at determining the factors that function downstream of FERONIA/LORELEI to decode species identity.

## Supporting information

Supplementary File Index

## Data availability statement

Raw data for the following figures is provided as listed. All original confocal time-lapse datasets and region of interest coordinates are available: https://doi.org/10.26300/b6za-nt53.

Figure 1-3 - Raw trace data provided in Supplementary File 1

Figure 1-3 – Raw statistical data provided in Supplementary File 2

## Code availability statement

All code used in this manuscript is supplied as supplementary data. Data for this study was collected using Zen Black (available through Zeiss), and the opensource ImageJ platform (available at https://imagej.nih.gov/ij). Ratiometric calculations were performed using a custom ImageJ .IJM file (Supplementary File 3) written by NDP that depends on the RatioPlus plugin (available at https://imagej.nih.gov/ij/plugins/ratio-plus.html). Data were analyzed using MATLAB (available at https://www.mathworks.com/products/matlab.html). The MATLAB script used to organize fluorescent data for plotting was written by NDP and is supplied as Supplementary File 4. The MATLAB script used to plot concatenated data as colored lines was written by NDP and is provided as Supplementary File 5. The MATLAB script used to plot concatenated data as boxplots was written by NDP and is supplied as Supplementary File 6. Supplementary File 6 relies on the ViolinPlot repository (available at https://github.com/bastibe/Violinplot-Matlab).

## Author contributions

NDP and MAJ designed and planned the experiments. NDP performed the experiments and collected the data. NDP analyzed the data. NDP and MAJ wrote the manuscript.

## Acknowledgements

We thank Ravishankar Palanivelu, Sharon Kessler, Jeff Harper, and Alexander Leydon for generating and/or providing transgenic lines for this study. We thank the Leduc Imaging Core Facility at Brown University and Geoff Williams for providing resources and expertise regarding time-lapse imaging. We thank Alison DeLong, Judith Bender, and Jenna Kotak for helpful discussions regarding this study. We thank Sherry Warner for maintaining plantings for this study. Funding for this project was provided by US National Science Foundation grant IOS-1353798 (MJ) and US National Institutes of Health Training Grant #T32-GM007601.

## Supplementary Figures

**Supplementary Figure 1.**
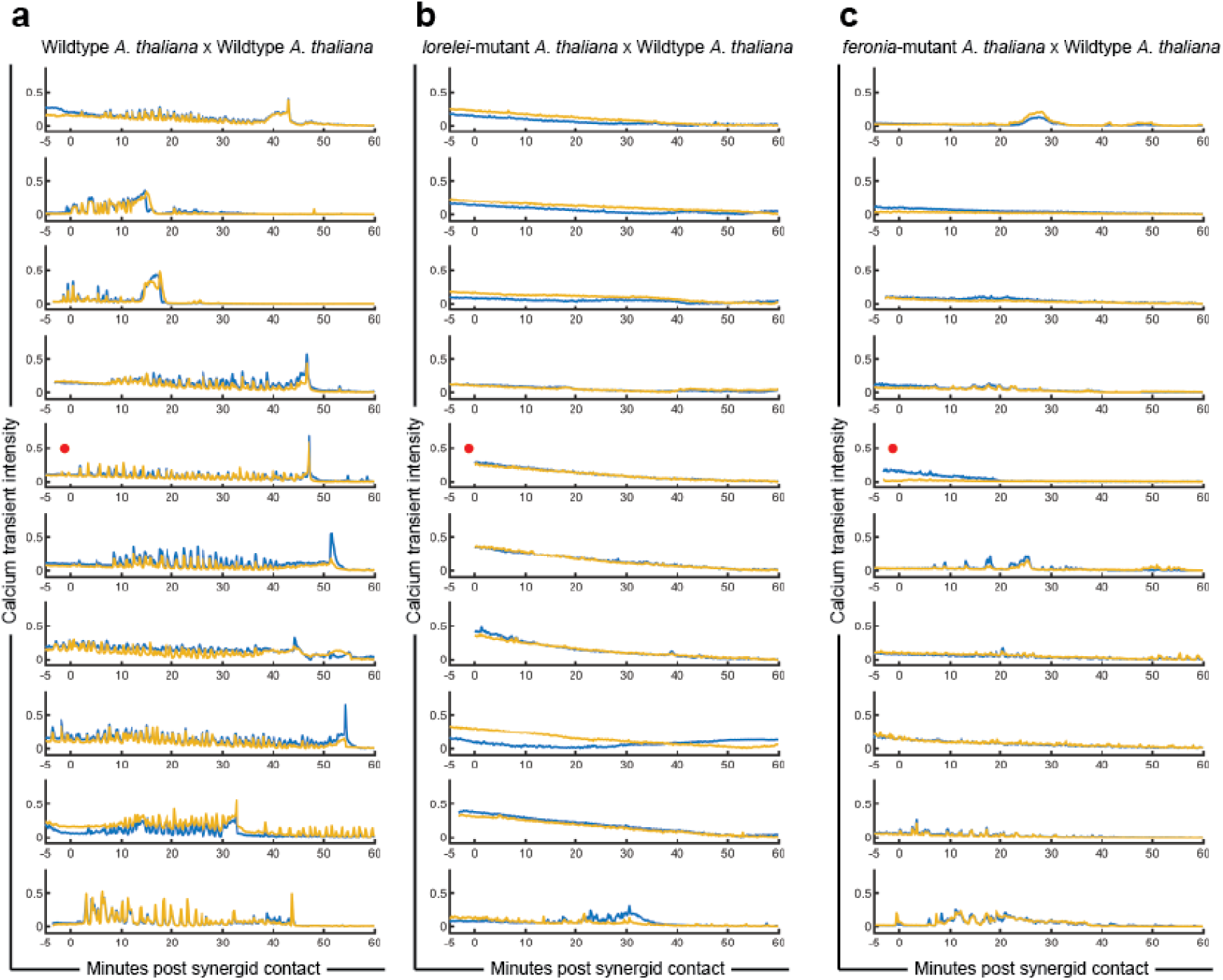
**a.** 10 replicate time-lapse calcium ion traces of pMYB98:YC3.60 in Wildtype *Arabidopsis thaliana* crossed as female with Wildtype *Arabidopsis thaliana* pollen. **b.** 10 replicate time-lapse calcium ion traces of pMYB98:YC3.60 in *lorelei*-mutant background *Arabidopsis thaliana* crossed as female with Wildtype *Arabidopsis thaliana* pollen. **c.** 10 replicate time-lapse calcium ion traces of pMYB98:YC3.60 in *feronia*-mutant background *Arabidopsis thaliana* crossed as female with Wildtype *Arabidopsis thaliana* pollen. For each trace left synergid region of interest plotted with blue line and right synergid region of interest plotted with yellow line. Traces shown in Figure 1 highlighted with red dot at left.

**Supplementary Figure 2.**
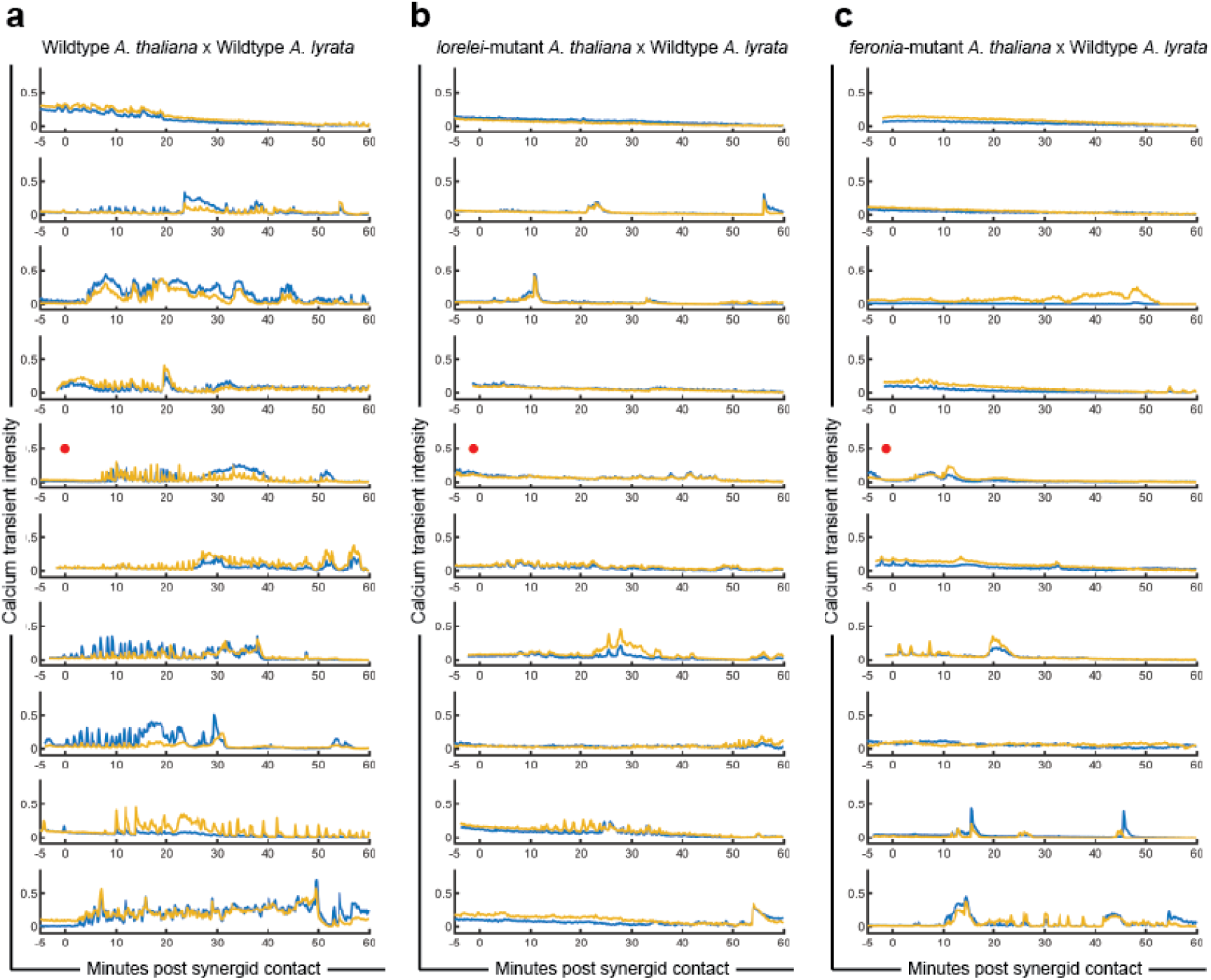
**a.** 10 replicate time-lapse calcium ion traces of pMYB98:YC3.60 in Wildtype *Arabidopsis thaliana* crossed as female with Wildtype *Arabidopsis lyrata* pollen. **b.** 10 replicate time-lapse calcium ion traces of pMYB98:YC3.60 in *lorelei*-mutant background *Arabidopsis thaliana* crossed as female with Wildtype *Arabidopsis lyrata* pollen. **c.** 10 replicate time-lapse calcium ion traces of pMYB98:YC3.60 in *lorelei-*mutant background *Arabidopsis thaliana* crossed as female with Wildtype *Arabidopsis lyrata* pollen. For each trace left synergid region of interest plotted with blue line and right synergid region of interest plotted with yellow line. Traces shown in Figure 2 highlighted with red dot at left.

**Supplementary Figure 3.**
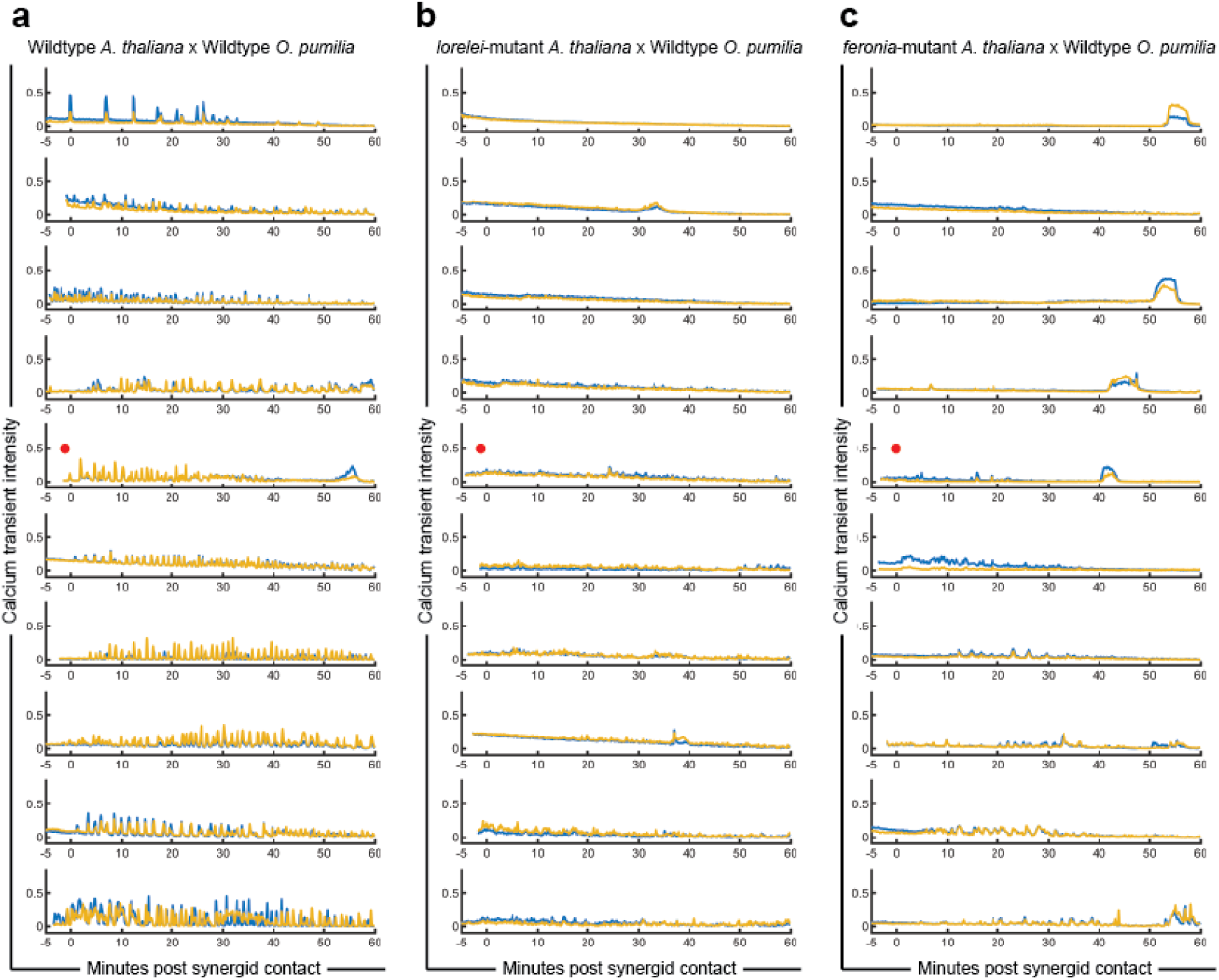
**a.** 10 replicate time-lapse calcium ion traces of pMYB98:YC3.60 in Wildtype *Arabidopsis thaliana* crossed as female with Wildtype *Olimarabidopsis pumilia* pollen. **b.** 10 replicate time-lapse calcium ion traces of pMYB98:YC3.60 in *lorelei*-mutant background *Arabidopsis thaliana* crossed as female with Wildtype *Olimarabidopsis pumilia* pollen. **c.** 10 replicate time-lapse calcium ion traces of pMYB98:YC3.60 in *feronia*-mutant background *Arabidopsis thaliana* crossed as female with Wildtype *Olimarabidopsis pumilia* pollen. For each trace left synergid region of interest plotted with blue line and right synergid region of interest plotted with yellow line. Traces shown in Figure 2 highlighted with red dot at left.

**Supplementary Figure 4.**
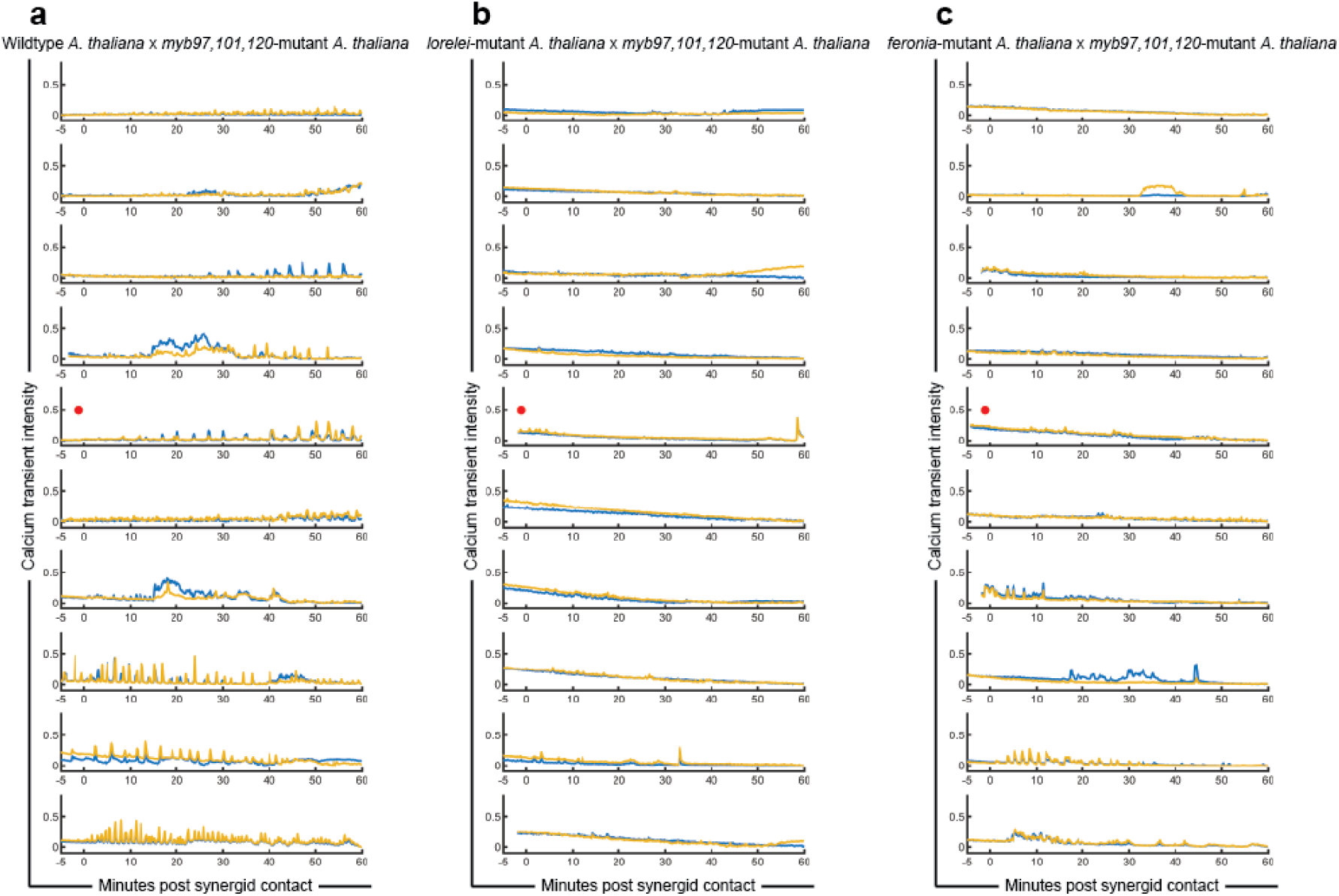
**a.** 10 replicate time-lapse calcium ion traces of pMYB98:YC3.60 in Wildtype *Arabidopsis thaliana* crossed as female with *myb97,101,120* triple-mutant *Arabidopsis thaliana* pollen. **b.** 10 replicate time-lapse calcium ion traces of pMYB98:YC3.60 in *lorelei*-mutant background *Arabidopsis thaliana* crossed as female with *myb97,101,120* triple-mutant *Arabidopsis thaliana* pollen. **c.** 10 replicate time-lapse calcium ion traces of pMYB98:YC3.60 in *feronia*-mutant background *Arabidopsis thaliana* crossed as female with *myb97,101,120* triple-mutant *Arabidopsis thaliana* pollen. For each trace left synergid region of interest plotted with blue line and right synergid region of interest plotted with yellow line. Traces shown in Figure 3 highlighted with red dot at left.

**Supplementary Figure 5.**
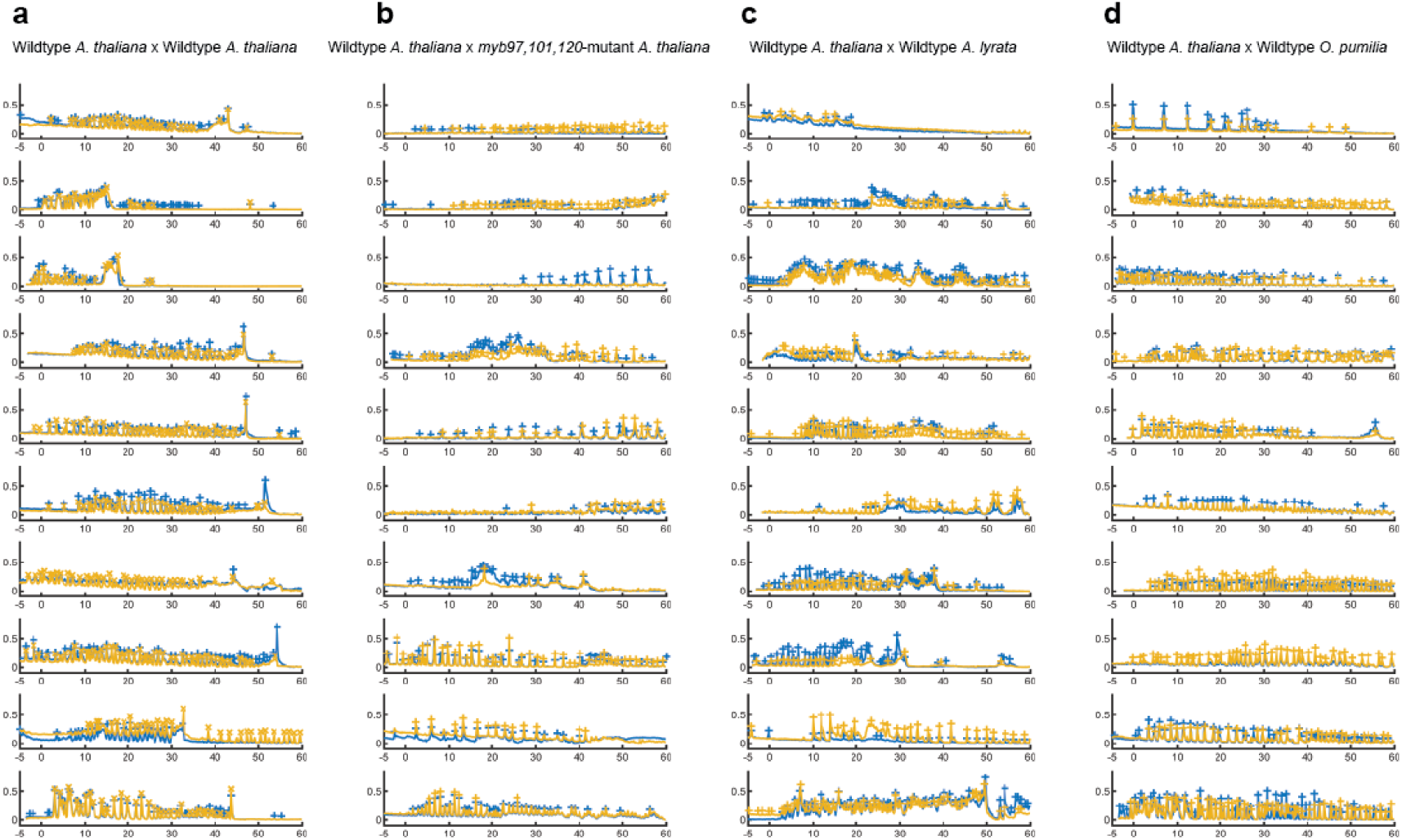
**a.** 10 replicate time-lapse calcium ion calcium ion traces of pMYB98:YC3.60 in Wildtype *Arabidopsis thaliana* crossed as female with Wildtype *Arabidopsis thaliana* pollen. **b.** 10 replicate time-lapse calcium ion traces of pMYB98:YC3.60 in Wildtype *Arabidopsis thaliana* crossed as female with *myb97,101,120* triple-mutant *Arabidopsis thaliana* pollen. **c.** 10 replicate time-lapse calcium ion traces of pMYB98:YC3.60 in Wildtype *Arabidopsis thaliana* crossed as female with Wildtype *Arabidopsis lyrata* pollen. For each trace left synergid region of interest plotted with blue line and right synergid region of interest plotted with yellow line. **d.** 10 replicate time-lapse calcium ion traces of pMYB98:YC3.60 in Wildtype *Arabidopsis thaliana* crossed as female with Wildtype *Olimarabidopsis pumilia* pollen. For each trace calcium ion peaks that were used for frequency analysis (with a prominence greater than 5x minimum prominence observed) are marked with ‘+’ for left synergid ROI and ‘x’ for right synergid ROI).

## Materials and Methods

### Transgenic Lines

Wildtype and *lorelei*-mutant background pMYB98:YC3.60^40^ lines were supplied by Dr. Ravishankar Palanivelu’s Laboratory at the University of Arizona. pMYB98:YC3.60 was introgressed into the *nta-1* mutant background and the *fer-4* mutant background which were gifted by Dr. Sharon Kessler’s Laboratory at Purdue University. The *Arabidopsis thaliana* pLAT52:dsRed line was gifted by Dr. Jeffery Harper’s Laboratory at The University of Nevada, Reno. *Olimarabidopsis pumilia* was transformed with pLAT52:dsRed using Agrobacterium-mediated transformation by Dr. Alexander Leydon (Current address: Department of Biology; University of Washington, Seattle). The *myb97, 101, 120* was generated by Dr. Alexander Leydon. All genotyping primer sequences used for this study are available upon request.

### Arabidopsis growth, pistil dissection, and time-lapse microscopy

Arabidopsis plants were grown under 16-hours-light, 8-hours-dark until mature. Pistils were emasculated 24 hours before dissections. Pistils were pollinated and left for 45 minutes before dissection. 200 microliters of Arabidopsis pollen growth media (5mM KCl, 0.01% H3BO3, 5mM CaCl2, 1mM MgSO4, 10% Sucrose, 1.5% NuSeive GTG Agarose, pH 7.6) was allowed to cool as a pad in a MatTek 35mm glass bottom dish with No. 1.5 coverglass (part no. P35G-1.5-20-C, https://www.mattek.com/store/p35g-1-5-20-c-case/). After media had cooled, pistils were dissected onto media and left in a humidity chamber at 22°CC for 5 hours. Dishes were imaged on a Zeiss LSM Laser Scanning Confocal Microscope at 20x optical zoom with 3.5x digital zoom. Images were taken with a 10 second interval starting before pollen tube contact with synergid cells and continuing for 1 hour after contact. For *Arabidopsis lyrata*, because we do not have a pollen-expressed fluorescent reporter in this species, we could not track the growth of the pollen tube tip after it had grown through the micropyle. To address this, during wildtype *A. thaliana* pMYB98:YC3.60 x *A. lyrata* crosses, we used the initial calcium ion transient observed in synergid cells as the marker for contact between the pollen tube tip and synergid cell. During *lorelei* pMYB98:YC3.60 x *A. lyrata* crossesand *feronia* pMYB98:YC3.60 x *A. lyrata* crosses, we used deformation of the synergid cell as the marker for contact between the pollen tube tip and synergid cell, as in these mutant backgrounds calcium ion oscillations are largely abolished^4^. For crosses using *A. thaliana* or *O. pumilia* pollen donors, pollen-expressed red fluorescence allowed us to track the pollen tube as it grew into the embryo sac and first contact was defined visually.

### Time-lapse data set analysis

To analyze time-lapse datasets, we use a custom ImageJ (https://imagej.nih.gov/ij/) macro (supplied in Supplementary File 4 as a commented, editable .ijm file). Any time-lapse movies suffering from drift were corrected using the MultiStackReg plugin (http://bradbusse.net/sciencedownloads.html). Supplementary File 4 splits multi-channel time-lapse datasets into constituent channels, asks the user to draw regions of interest around the left and right synergid cells, subtracts any pollen-expressed fluorescence from synergid cell channels, and uses the RatioPlus plugin (https://imagej.nih.gov/ij/plugins/ratio-plus.html) to calculate relative FRET efficiency within the regions of interest. Following this, we remove bright outliers and despeckle the output to remove noise resulting from our data collection process. The ratiometric results are saved automatically as .csv files, along with the summed projection over which the user draws the regions of interest around the synergid cells, the coordinates of regions of interest themselves, and a flattened composite time-lapse to the directory from which the time-lapse dataset is opened.

### FRET data analysis

Data was analyzed using MATLAB (available at https://www.mathworks.com/products/matlab.html). The MATLAB script used to concatenate fluorescent data (Supplementary File 1) was written by NDP and is supplied as Supplementary File 5. The MATLAB script used to plot concatenated data as colored lines was written by NDP and is provided as Supplementary File 6. The MATLAB script used to plot concatenated data as boxplots was written by NDP and is supplied as Supplementary File 7. Supplementary File 7 relies on the ViolinPlot repository (available at https://github.com/bastibe/Violinplot-Matlab). For Figures 1-3, calcium ion peaks were analyzed if they were greater than 1.1x the minimum prominence value registered for their respective trace. This cutoff value was chosen such that noise could be excluded from the analysis while still allowing traces with few/small prominences (those onto *lorelei* or *feronia* females) could still be analyzed. For Figure 4, calcium ion peaks with prominences greater than 5x the minimum prominence value registered for their respective traces were analyzed.

### Data plotting and figure construction

For each cross scheme all ten replicates were sorted according to average calcium peak prominence. Traces shown in Figures 1-3 were chosen as they are the trace with median peak prominence. For each statistical plots in Figures 1-4 median value denoted by red line. 95% confidence interval about the median is denoted by notches. Box defines interquartile range. Length of whiskers is calculated as 1.5*IQR. ANOVA and post-hoc multiple comparison of means supplied as Supplementary File 2 and Supplementary File 3.

