## Supplementary material for "Synergid calcium ion oscillations define a new feature of pollen tube reception critical for blocking interspecific hybridization": Supplementary_File_04.ijm

```
run("Set Measurements...", "mean redirect=None decimal=3");
//get current directory
dir = getDirectory("image");
//get current title
curtitle = getTitle();
//split stack into channels
run("Split Channels");
//make titles for each channel
CFP_title = "C1-" + curtitle;
FRET_title = "C2-" + curtitle;
YFP_title = "C3-" + curtitle;
Cy5_title = "C5-" + curtitle;
PT_title = "C4-" + curtitle;
//get projection from FRET channel
selectWindow(FRET_title);
run("Z Project...", "projection=[Max Intensity]");
//ask user for PT left synergid ROI
setTool("polygon");
waitForUser("Please create a selection to indicate\nthe PTleft synergid, then\nclick OK to proceed.");
roiManager("Add");
//save left synergid coordinates
roiManager("Select", 0);
getSelectionCoordinates(x, y);
for (i=0; i<x.length; i++)
	print(i+" "+x[i]+" "+y[i]);
selectWindow("Log");
saveAs("Text", dir + "PTleft_synROI_coordinates");
print("\\Clear");
selectWindow("Log"); 
run("Close"); 
//ask user for PT right synergid ROI
waitForUser("Please create a selection to indicate\nthe PTright synergid, then\nclick OK to proceed.");
roiManager("Add");
//save right synergid coordinates
roiManager("Select", 1);
getSelectionCoordinates(x, y);
for (i=0; i<x.length; i++)
	print(i+" "+x[i]+" "+y[i]);
selectWindow("Log");
saveAs("Text", dir + "PTright_synROI_coordinates");
selectWindow("Log"); 
run("Close"); 
//save projection
selectWindow("MAX_" + FRET_title);
saveAs("Tiff");
//subtract any pollen fluorescence from CFP
imageCalculator("Subtract create 32-bit stack", CFP_title, PT_title);
imageCalculator("Add create 32-bit stack", "Result of C1-" + curtitle, YFP_title);
//subtract any pollen fluorescence from FRET
imageCalculator("Subtract create 32-bit stack", FRET_title, PT_title);
//take ratio of (FRET-RFP)/(CFP-RFP)
run("Ratio Plus", "image1=[Result of C2-" + curtitle + "] image2=[Result of C1-" + curtitle + "] background1=10 clipping_value1=0 background2=10 clipping_value2=0 multiplication=10");
//despeckle ratio
run("Despeckle", "stack");
//remove bright noise from ratio
run("Remove Outliers...", "radius=10 threshold=10 which=Bright stack");
//enhance ratio contrast
run("Enhance Contrast...", "saturated=0.03 normalize process_all use");
//measure pulses
selectWindow("Ratio");
roiManager("Select", newArray(0,1));
roiManager("Multi Measure");
saveAs("Results");
roiManager("Delete");
//select look up table
selectWindow("Ratio");
run("mpl-viridis");
//make Cy5 32bit
selectWindow("C4-" + curtitle);
run("32-bit");
//make PT 32bit
selectWindow("C5-" + curtitle);
run("32-bit");
//make PT+ratio+Cy5 composite
run("Merge Channels...", "c1=Ratio c2=C4-" + curtitle + " c3=C5-" + curtitle + " create keep");
run("Flatten");
//save composite
selectWindow("Composite");
saveAs("Tiff");
```
