## Supplementary material for "Synergid calcium ion oscillations define a new feature of pollen tube reception critical for blocking interspecific hybridization": Supplementary_File_05.m

```
prom_cut = 5
%define datasets
% names = {WT_Athaliana_female_X_WT_Athaliana_male,...
%     lorelei_Athaliana_female_X_WT_Athaliana_male,...
%     feronia_Athaliana_female_X_WT_Athaliana_male,...
%     WT_Athaliana_female_X_myb_Athaliana_male,...
%     lorelei_Athaliana_female_X_myb_Athaliana_male,...
%     feronia_Athaliana_female_X_myb_Athaliana_male,...
%     WT_Athaliana_female_X_WT_Alyrata_male,...
%     lorelei_Athaliana_female_X_WT_Alyrata_male,...
%     feronia_Athaliana_female_X_WT_Alyrata_male,...
%     WT_Athaliana_female_X_WT_Opumilia_male,...
%     lorelei_Athaliana_female_X_WT_Opumilia_male,...
%     feronia_Athaliana_female_X_WT_Opumilia_male,...
%     }
names = {WT_Athaliana_female_X_WT_Athaliana_male,...
    WT_Athaliana_female_X_myb_Athaliana_male,...
    WT_Athaliana_female_X_WT_Alyrata_male,...
    WT_Athaliana_female_X_WT_Opumilia_male,...
    }
%for each dataset
for i = 1:1:numel(names)
%     figure
    %for each replicate in that datase
    for j = 1:1:numel(names{i})
        %left synergid data is 1st column
        left = table2array(names{i}{j}(:,1));
        %take data after point of first contact
        if length(left) > 399 
            left = left(length(left)-399:length(left));
        else
            left = left;
        end
        %make lowest value set to 0
        left = left-min(left);
        %concatenate left synergid data into new array
        ldata{i}{j} = left;
        %repeat above steps for right synergid data
        right = table2array(names{i}{j}(:,2));
        if length(right) > 399
            right = right(length(right)-399:length(right));
        else
            right = right;
        end
        right = right-min(right);
        rdata{i}{j} = right;       
        %find local maxima in left synergid data
        [lpks{i}{j}(:,1),llocs{i}{j}(:,1),lwidths{i}{j}(:,1),lproms{i}{j}(:,1)] = findpeaks(left,'MinPeakProminence',(prom_cut*min(findpeaks(left))));
        %find frequency of local maxima
        ldiff{i}{j} = [0;diff(llocs{i}{j}(:,1))];
%         leftline = prom_cut*min(findpeaks(left));
%         subplot(10,1,j)
%         hold on
%         findpeaks(left,'MinPeakProminence',(prom_cut*min(findpeaks(left))))
%         line([0 399], [leftline leftline],'Color','blue')
        %repeat for right synergid data
        [rpks{i}{j}(:,1),rlocs{i}{j}(:,1),rwidths{i}{j}(:,1),rproms{i}{j}(:,1)] = findpeaks(right,'MinPeakProminence',(prom_cut*min(findpeaks(right))));
%         rightline = prom_cut*min(findpeaks(right));
        rdiff{i}{j} = [0;diff(rlocs{i}{j}(:,1))];
%         findpeaks(right,'MinPeakProminence',(prom_cut*min(findpeaks(right))))
%         line([0 399], [rightline rightline],'Color','red')
    end
end
```
