## Supplementary material for "Synergid calcium ion oscillations define a new feature of pollen tube reception critical for blocking interspecific hybridization": Supplementary_File_06.m

```
% % datasets: 1=wt x wt, 2 = lre x wt, 3 = fer x wt
% % 4 = wt x myb, 5 = lre x myb, 6 = fer x myb,
% % 7 = wt x alyr, 8 = lre x alyr, 9 = fer x alyr, 
% % 10 = wt x opumil, 11 = lre x opumil, 12 = fer x opumil

%define dataset or range of datasets as variable b
for b = 1:1:4;
    %make figure
    figure
    %make subplot for each replicate
    for i = 1:1:size(ldata{1,b},2)
        subplot(size(ldata{1,b},2),1,i)
        %define left synergid as first column in replicate
        left = ldata{1,b}{1,wtlist{1,b}(i)};
        leftx = llocs{1,b}{1,wtlist{1,b}(i)};
        lefty = lpks{1,b}{1,wtlist{1,b}(i)};
        %define right synergid as second column in replicate
        right = rdata{1,b}{1,wtlist{1,b}(i)};
        rightx = rlocs{1,b}{1,wtlist{1,b}(i)};
        righty = rpks{1,b}{1,wtlist{1,b}(i)};
        %plot left data
        hold on
%         plot(flipud(left),'Color',[0, 0.4470, 0.7410])
%         plot((size(left)+1)-leftx,lefty+0.05,'+','Color',[0, 0.4470, 0.7410])
        %plot right data
        plot(flipud(right),'Color',[0.9290, 0.6940, 0.1250])
        plot((size(right)+1)-rightx,righty+0.05,'+','Color',[0.9290, 0.6940, 0.1250])
        %define box dimensions
        xlim([0 390])
        ylim([-.1 .85])
%         xticks([0,30,90,150,210,270,330,390])
%         xticklabels({'-5','0','10','20','30','40','50','60'})
        xticks([0,60,120,180,240,300,360,390])
        xticklabels({'60','50','40','30','20','10','0','-5'})
        yticks([0, 0.5, 1])
        yticklabels({'0','0.5','1'})
        box off
        view(180,270)
    end
end
```
