## Supplementary material for "Synergid calcium ion oscillations define a new feature of pollen tube reception critical for blocking interspecific hybridization": Supplementary_File_07.m

```
%define left synergid metrics
l_items = {lpks,llocs,lwidths,lproms,ldiff};
%define right synergid metrics
r_items = {rpks,rlocs,rwidths,rproms,rdiff};
%concatenate left and right metrics of every replicate or each crossing scheme into new catitems array
for i = 1:1:numel(l_items)
    for j = 1:1:numel(l_items{i})
        for k = 1:1:numel(l_items{i}{j})
        	cat_items{i}{j} = [l_items{i}{j}{k};r_items{i}{j}{k}];
        end
    end
end
%clear variables
%clear('i','j','k','l_items','r_items','ldiff','rdiff')
%define titles
titles = {'Height','Location','Width','Prominance','Difference'}
%define which metric or set of metrics you want to plot
for i = 5;
    %for each datapoint in that metric define a group for the plot
    for j = 1:1:numel(cat_items{i});
        cat_groups{i}{j} = ones(size(cat_items{i}{j},1),1)*j;
    end
    %define values as datapoints
    values{i} = vertcat(cat_items{i}{:});
    %define groups as group values
    groups{i} = vertcat(cat_groups{i}{:});
    %make a figure
    figure
    hold on
    %title the figure with appropriate title
    title(titles{i})
    %define value as appropriate values
    value = values{i};
    %define group as appropriate groups
    group = groups{i};
    %make violin plot from data
    violinplot(value,group,'ViolinAlpha', 0.05,'ViolinColor',[0 0 0],'ShowData',false);
    %overlay boxplot of data
    boxplot(value,group,'Notch','on','symbol','');
    %overlay scattered data points with jitter to make them visable
    jit = randn(1,length(group))*0.06;
    jit = jit(:);
    scatter(group+jit,value,20,'filled','k','MarkerFaceAlpha',.1);
    %separate groups with lines
    line([3.5 3.5],[-.5 .7],'color','k')
    line([6.5 6.5],[-.5 .7],'color','k')
    line([9.5 9.5],[-.5 .7],'color','k')
end
hold on
[~,~,stats] = anova1(value,group);
multcompare(stats,'CType','bonferroni');
view(270,90)
% clear('ans','b','cat_groups','cat_items','group','groups','i','j','jit',...
%     'ldata','left','llocs','lpks','lproms','lwidths','names','rdata','right',...
%     'rlocs','rpks','rproms','rwidths','titles','value','values','stats','c')
```
